# Energy conservation via hydrogen cycling in the methanogenic archaeon *Methanosarcina barkeri*

**DOI:** 10.1101/335794

**Authors:** Gargi Kulkarni, Thomas D. Mand, William W. Metcalf

## Abstract

Energy conservation via hydrogen cycling, which generates proton motive force by intracellular H_2_ production coupled to extracellular consumption, has been controversial since it was first proposed in 1981. It was hypothesized that the methanogenic archaeon *Methanosarcina barkeri* is capable of energy conservation via H_2_ cycling, based on genetic data that suggest H_2_ is a preferred, but non-essential, intermediate in the electron transport chain of this organism. Here, we characterize a series of hydrogenase mutants to provide direct evidence of H_2_ cycling. *M. barkeri* produces H_2_ during growth on methanol, a phenotype that is lost upon mutation of the cytoplasmic hydrogenase encoded by *frhADGB*, although low levels of H_2_, attributable to the Ech hydrogenase, accumulate during stationary phase. In contrast, mutations that conditionally inactivate the extracellular Vht hydrogenase are lethal when expression of the *vhtGACD* operon is repressed. Under these conditions H_2_ accumulates, with concomitant cessation of methane production and subsequent cell lysis, suggesting that the inability to recapture extracellular H_2_ is responsible for the lethal phenotype. Consistent with this interpretation, double mutants that lack both Vht and Frh are viable. Thus, when intracellular hydrogen production is abrogated, loss of extracellular H_2_ consumption is no longer lethal. The common occurrence of both intracellular and extracellular hydrogenases in anaerobic microorganisms suggests that this unusual mechanism of energy conservation may be widespread in nature.

**Importance:** Adenosine triphosphate (ATP) is required by all living organisms to facilitate essential endergonic reactions required for growth and maintenance. Although synthesis of ATP by substrate-level phosphorylation is widespread and significant, most ATP is made via the enzyme ATP synthase, which is energized by transmembrane chemiosmotic gradients. Therefore, establishing this gradient across the membrane is of central importance to sustaining life. Experimental validation of H_2_ cycling adds to a short list of mechanisms for generating a transmembrane electrochemical gradient that is likely to be widespread, especially among anaerobic microorganisms.

## Introduction

An essential requirement for life is the ability to couple exergonic metabolism to the endergonic synthesis of adenosine triphosphate (ATP). While some ATP is made by direct phosphorylation of adenosine diphosphate using “high-energy” metabolites such as phosphoenolpyruvate or 1,3-diphosphoglycerate, the vast majority is produced via the enzyme ATP synthase using energy stored in a transmembrane proton (or sodium) gradient. These electrochemical gradients are typically established during the process of electron transport by membrane proteins that couple exergonic redox reactions to generation of an ion-motive force by one of three general mechanisms: (*i*) vectorial proton pumping, (*ii*) scalar movement of protons across the membrane, as in the Q-cycle or Q-loop, or (*iii*) coupled reactions that consume protons within the cell and produce protons on the outside (1, 2). Given the importance of this process, it is not surprising that this central aspect of living systems has been the subject of intense study (and at least three Nobel Prizes). Indeed, we now possess a detailed, molecular-level understanding of chemiosmotic energy conservation as it applies to photosynthesis and aerobic respiration in a wide variety of organisms including eukaryotes, bacteria, and archaea. Nevertheless, unique and sometimes surprising mechanisms for generation of chemiosmotic gradients continue to be found, including sodium-pumping methyltransferases in methanogenic archaea (3), electrogenic formate:oxalate antiporters in bacteria (4, 5), and light-driven, proton-pumping rhodopsins (6).

A controversial, and as yet unproven, mechanism for creating transmembrane proton gradients called H_2_ cycling was proposed by Odom and Peck in 1981 to explain ATP synthesis in sulfate-reducing bacteria (7). In this proposed energy-conserving process, protons in the cytosol are reduced to molecular H_2_ by enzymes known as hydrogenases. The H_2_ so produced then diffuses across the membrane where it is re-oxidized by extracellular hydrogenases, releasing protons that contribute to a transmembrane proton gradient that can be used to make ATP. The electrons produced by this reaction are returned to the cytoplasm via a membrane-bound electron transport chain, completing the redox process.

Although H_2_ cycling has been suggested to occur in a number of anaerobic organisms (7-11), the hydrogen cycling hypothesis has not been widely accepted. A key argument against the idea is based on the high diffusion rate of molecular hydrogen. Thus, unless extracellular recapture is exceptionally efficient, hydrogen produced in the cytoplasm would be easily lost, resulting in redox imbalance and presumably cell death. Nevertheless, experimental demonstration of simultaneous production and consumption of H_2_ by *Desulfovibrio* supports the model (12), as does metabolic modeling (13). However, other data are inconsistent with the idea, including the ability of hydrogenase mutants to grow on lactate (14) and the inability of high external H_2_ pressures to inhibit substrate catabolism (15). Thus, the H_2_ cycling model for energy conservation remains unproven.

Based on series of genetic experiments, we proposed that the methanogenic archaeon, *Methanosarcina barkeri*, employs H_2_ cycling during growth on one-carbon (C-1) substrates and acetate (16, 17). During growth on C-1 compounds such as methanol, the putative cycling pathway would produce H_2_ using the cytoplasmic F420-dependent (Frh) and energy-converting ferredoxin-dependent (Ech) hydrogenases, while H_2_ production during growth on acetate would be mediated solely by Ech. Both pathways would converge on the methanophenazine-dependent hydrogenase (Vht), which is thought to have an active site on the outer face of the cell membrane (18), to consume extracellular H_2_ and deliver electrons to the membrane-bound electron transport chain, where they serve to reduce the coenzyme M-coenzyme B heterodisulfide (CoM-S-S-CoB) produced during the production of methane (Fig 1). However, these genetic studies remain incomplete because neither the role of Vht, nor the production and consumption of hydrogen were examined. Here we explicitly test both, providing strong experimental support for the role of H_2_ cycling in energy conservation in *M. barkeri*.

**Figure 1.**
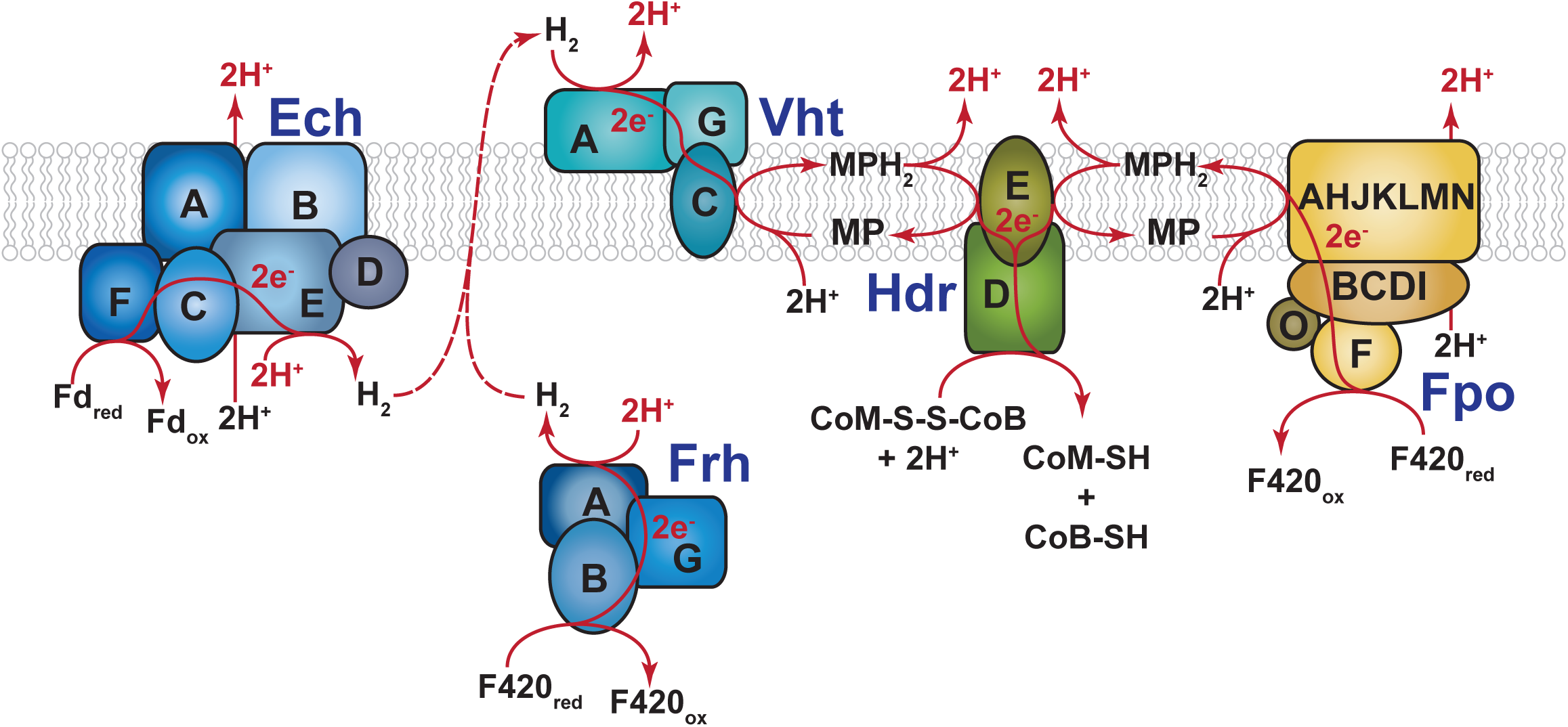
The putative H_2_ cycling electron transport chain of *M. barkeri*. Growth on C-1 substrates generates reduced cofactor F420 (F420_red_), which is a hydride carrying cofactor analogous to NADH, and the reduced form of the small electron-carrying protein ferredoxin (Fd_red_). During aceticlastic methaogensis only Fd_red_ is produced. These reduced electron carriers are re-oxidized in the cytoplasm by the Frh and Ech hydrogenases, respectively, with concomitant consumption of protons to produce molecular H_2_. H_2_ subsequently diffuses out of the cell where it is re-oxidized by the Vht hydrogenase, which has an active site located on the outer face of the cell membrane. This reaction releases protons on the outside of the cell and produces reduced methanophenazine (MPH_2_), a membrane-bound electron carrier analogous to ubiquinone. MPH_2_ subsequently delivers electrons to the enzyme heterodisulfide reductase (Hdr), which serves as the terminal step in the *Methanosarcina* electron transport chain. This final reaction regenerates coenzyme B (CoB-SH) and coenzyme M (CoM-SH) from the mixed disulfide (CoM-S-S-CoB), which is produced from the free thiol cofactors during methanogenic metabolism. Electron (e^-^) flow and scalar protons (H^+^) are shown in red. It should be noted that *M. barkeri* can also re-oxidize F420_red_ using the membrane-bound, proton-pumping F420-dehydrogenase (Fpo). Thus, the cell has a branched electron transport chain and, therefore, is not dependent on H_2_ cycling during growth on methylotrophic substrates (16); however, both pathways for electron transport from F420 have identical levels of energy conservation: namely 4 H^+^/2e^-^. It should also be noted that the Ech hydrogenase acts as a proton pump in addition to its role in H_2_ cycling, thus electron transport during aceticlastic methanogenesis conserves 6H^+^/2e^-^. Individual subunits of the various enzymes are indicated by letters (*e.g.* A, B, C…).

## Results and Discussion

### Hydrogenases of *M. barkeri*

Three distinct types of hydrogenases are encoded by *M. barkeri* Fusaro (Fig. S1) (19). The F420-reducing hydrogenase (Frh) is a cytoplasmic, 3-subunit (α, β, and γ) enzyme encoded by the *frhADGB* operon, which also includes a maturation protease, FrhD (20). This enzyme couples the oxidation/reduction of the deazaflavin cofactor F420 with production/consumption of H_2_. The membrane-bound Vht hydrogenase utilizes the quinone-like electron carrier, methanophenazine, as a cofactor (21). Like Frh, Vht is a 3-subunit enzyme encoded by a four-gene operon (*vhtGACD*) that includes a maturation protease, VhtD (19). *M. barkeri* also encodes homologs of both the *frh* and *vht* operons (the *freAEGB* and *vhxGAC* operons, respectively); however, multiple lines of evidence suggest these genes are incapable of producing active hydrogenases (16, 22). Thus, the presence of these genes has no bearing on the results presented herein. The final hydrogenase encoded by *M. barkeri* is a membrane-bound, <u>e</u>nergy-<u>c</u>onverting <u>h</u>ydrogenase (Ech), which couples the oxidation/reduction of ferredoxin and H_2_ to the production/consumption of a proton motive force (23, 24). Thus, the enzyme can use proton motive force to drive the endergonic reduction of ferredoxin by H_2_, which is required for CO_2_ reduction during hydrogenotrophic methanogenesis and for biosynthesis during growth by H_2_-dependent reduction of C-1 compounds (methyl-reducing methanogenesis). During both methylotrophic and aceticlastic methanogenesis, Ech is believed to couple oxidation of reduced ferredoxin to production of proton motive force and H_2_. The hydrogen thus produced would need to be recaptured by Vht in a putative H_2_ cycling process that contributes to proton motive force (Fig 1) (17).

### The cytoplasmic Frh hydrogenase is responsible for production of H_2_ during growth on methanol

A number of studies have shown that assorted *Methanosarcina* strains produce H_2_ during growth on methylotrophic and aceticlastic substrates (9, 25-30); however, to our knowledge this has never been assessed in *M. barkeri* strain Fusaro. To test this, we quantified the accumulation of CH_4_ and H_2_ during growth on methanol medium (Fig 2). Consistent with the hydrogen-cycling hypothesis, we observed significant H_2_ production, which reached a maximum partial pressure of *ca.* 20 Pa near the end of exponential growth. As expected, the culture also produced substantial levels of methane. As previously observed (16), a mutant lacking Frh (WWM115, Table S1) grew at a slower rate than its isogenic parent, and produced somewhat smaller amounts of methane. Very little H_2_ (< 4 Pa) was produced during growth of the Δ*frh* mutant; however, after growth ceased, the H_2_ concentration slowly rose, reaching a maximum level of 7 Pa. Thus, Frh is responsible for most hydrogen production during growth of *M. barkeri* Fusaro on methanol, although, some hydrogen is still produced in the Δ*frh* mutant. As will be shown below, Ech is probably responsible for the low levels of H_2_ seen in the Δ*frh* mutant.

**Figure 2.**
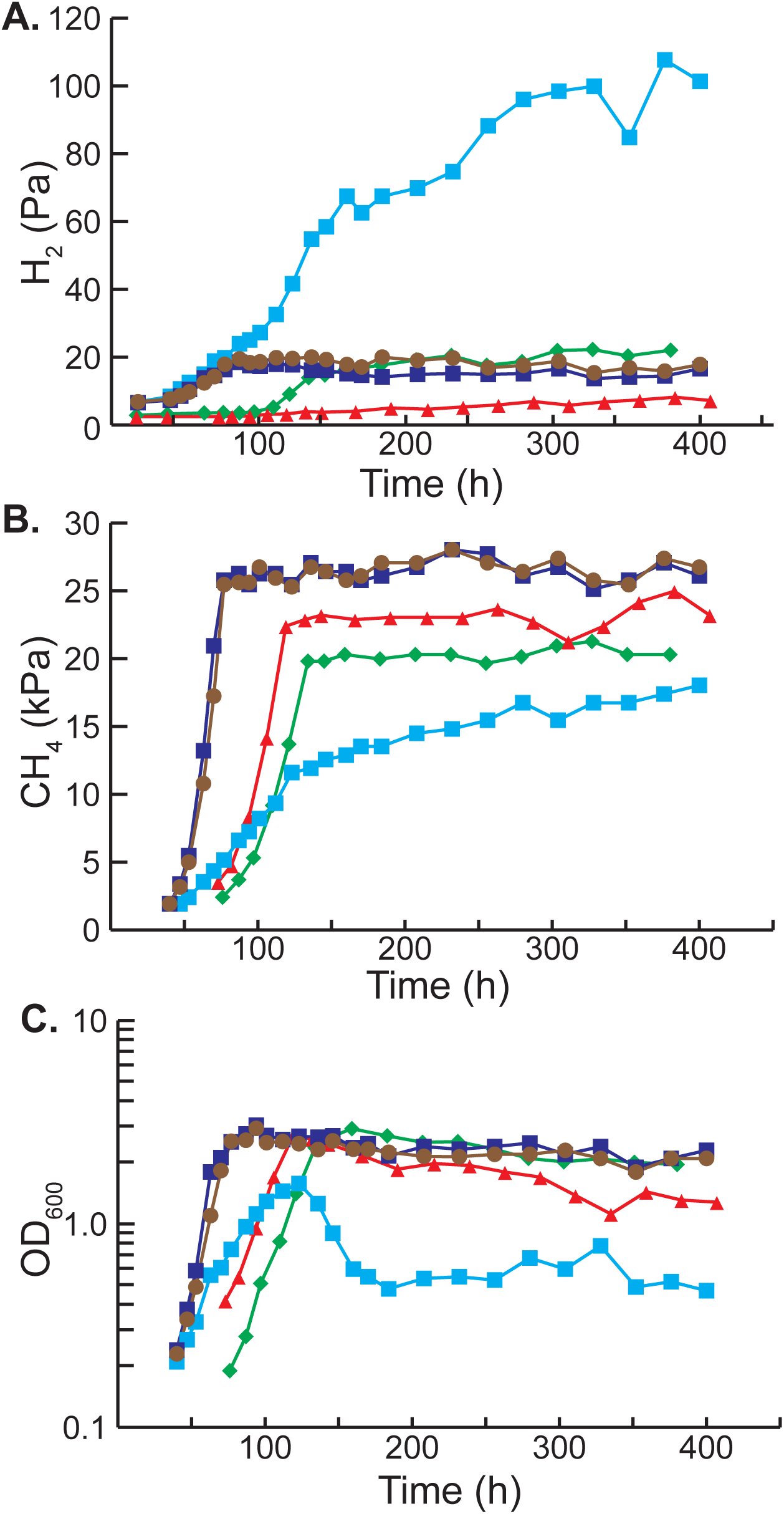
Hydrogen and methane production during methylotrophic growth. The partial pressures of H_2_ (*panel A*) and methane (*panel B*) were monitored during the course of growth (as indicated by optical density, *panel C*) in methanol medium for various *M. barkeri* strains. Strains used were: *M. barkeri* isogenic parental strain (brown circles, WWM85), tetracycline-regulated *vht* mutant (WWM157) with tetracycline (dark blue squares) and without tetracycline (light blue squares), *frh* deletion mutant (red triangles, WWM115), and *frh/vht* double deletion mutant (green diamonds, WWM351). Measurements were performed in triplicates as described in the methods section. Complete strain genotypes can be found in Table S1.

### Vht activity is required for viability of *M. barkeri*

To investigate the role of Vht during growth of *M. barkeri*, we attempted to delete the *vhtGACD* operon via homologous gene replacement (31, 32). However, despite numerous attempts, including selection on a variety of media, with and without supplementation of potential biosynthetic intermediates, no mutant colonies were obtained. We also attempted to delete the *vht* operon using the markerless deletion method of genetic exchange (33). This method relies on construction of a merodiploid strain with both mutant and wild type alleles. Upon segregation of the merodiploid, 50% of the recombinants are expected to be mutants if there is no selective pressure against the mutant allele. However, if the mutation causes a reduction in growth rate (with lethality being the most extreme case), the probability of obtaining recombinants with the mutant allele is severely reduced. We tested 101 haploid recombinants obtained from a *vhtGACD*^+^*/ΔvhtGACD* merodiploid; all carried the wild-type *vht* allele. Taken together, these data suggest that the *vhtGACD* operon is critical for normal growth of *M. barkeri*.

To test whether Vht is essential, we constructed a mutant in which the *vht* operon was placed under control of a tightly regulated, tetracycline-dependent promoter (34). We then examined the viability of the mutant and its isogenic parent by spotting serial dilutions on a variety of media, with and without tetracycline. As shown in Figure 3, the P_*tet*_*∷vht* mutant is unable to grow in the absence of the inducer, but grew well when tetracycline was added, whereas the isogenic parent grew with or without the addition of tetracycline. These phenotypes were observed on a variety of media including (*i*) methanol, (*ii*) methanol plus H_2_, (*iii*) H_2_/CO_2_, and (*iv*) acetate, which were chosen because they encompass growth conditions that require each of the four known methanogenic pathways used by *M. barkeri* (Fig 4). It should be stressed that the P_*tet*_*∷vht* mutant used in this experiment was pre-grown in the presence of inducer. Thus, at the start of the experiment, all cells have active Vht. However, during cultivation in the absence of tetracycline, pre-existing Vht is depleted by protein turnover and cell division, thereby allowing characterization of the Vht-deficient phenotype. The absence of growth of the diluted cultures in all media shows that Vht is essential for growth via the methylotrophic (methanol), methyl-reducing (methanol plus H_2_), hydrogenotrophic (H_2_/CO_2_) and aceticlastic (acetate) methanogenic pathways.

**Figure 3.**
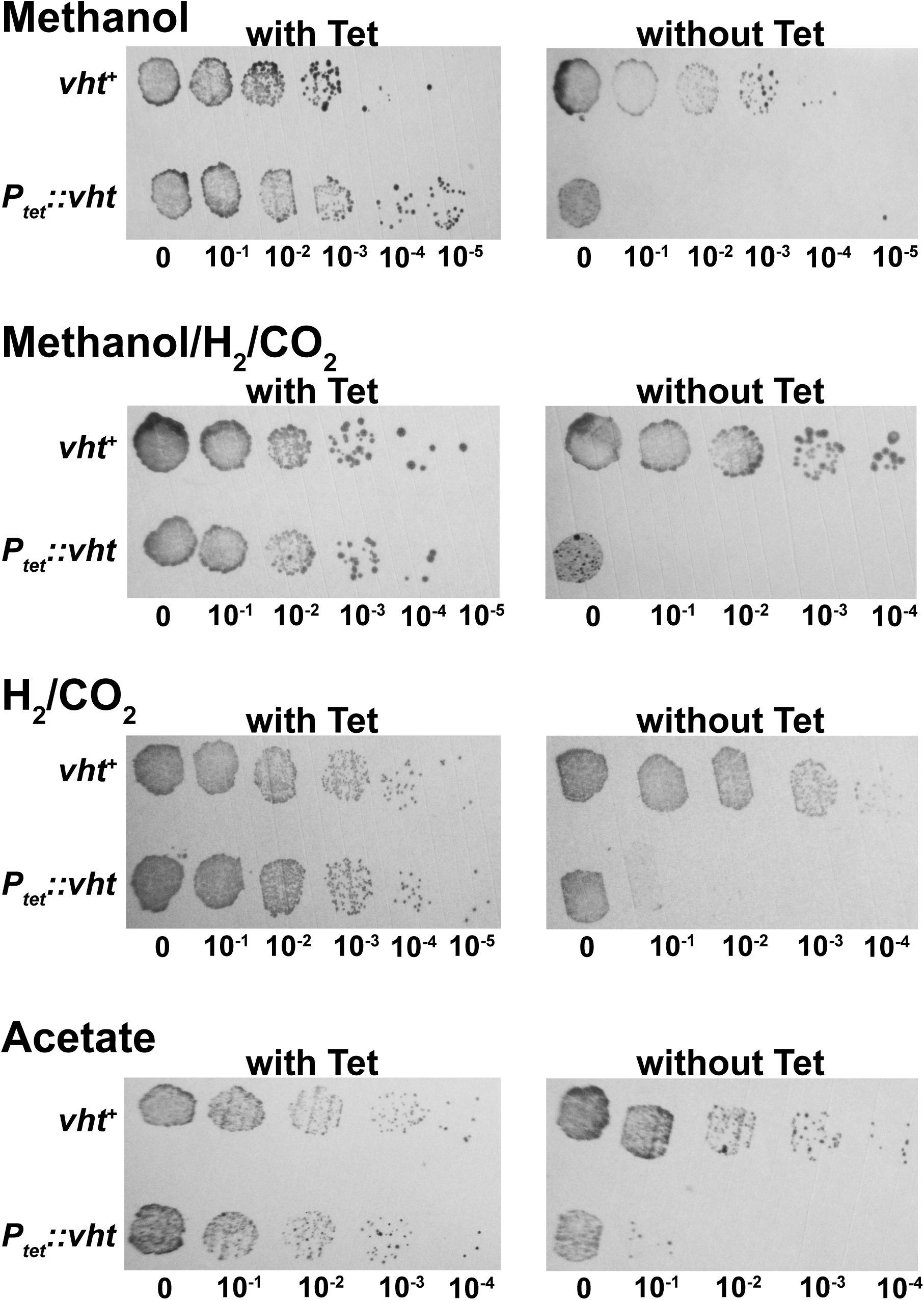
Essentiality of the Vht hydrogenase in *M. barkeri*. Cultures of the P_*tet*_*∷vht* mutant (WWM157) and its isogenic parent (WWM154) were adapted to four different substrates of interest (and in the presence of tetracycline for P_*tet*_*∷vht*), then washed, serially diluted, and incubated with each substrate with and without tetracycline (Tet). The media used indicate the ability to grow via each of the four known methanogenic pathways: (i) methylotrophic (methanol), (ii) methyl-reduction (methanol/H_2_/CO2), (iii) hydrogenotrophic (H_2_/CO_2_), and (iv) aceticlastic (acetate).

**Figure 4.**
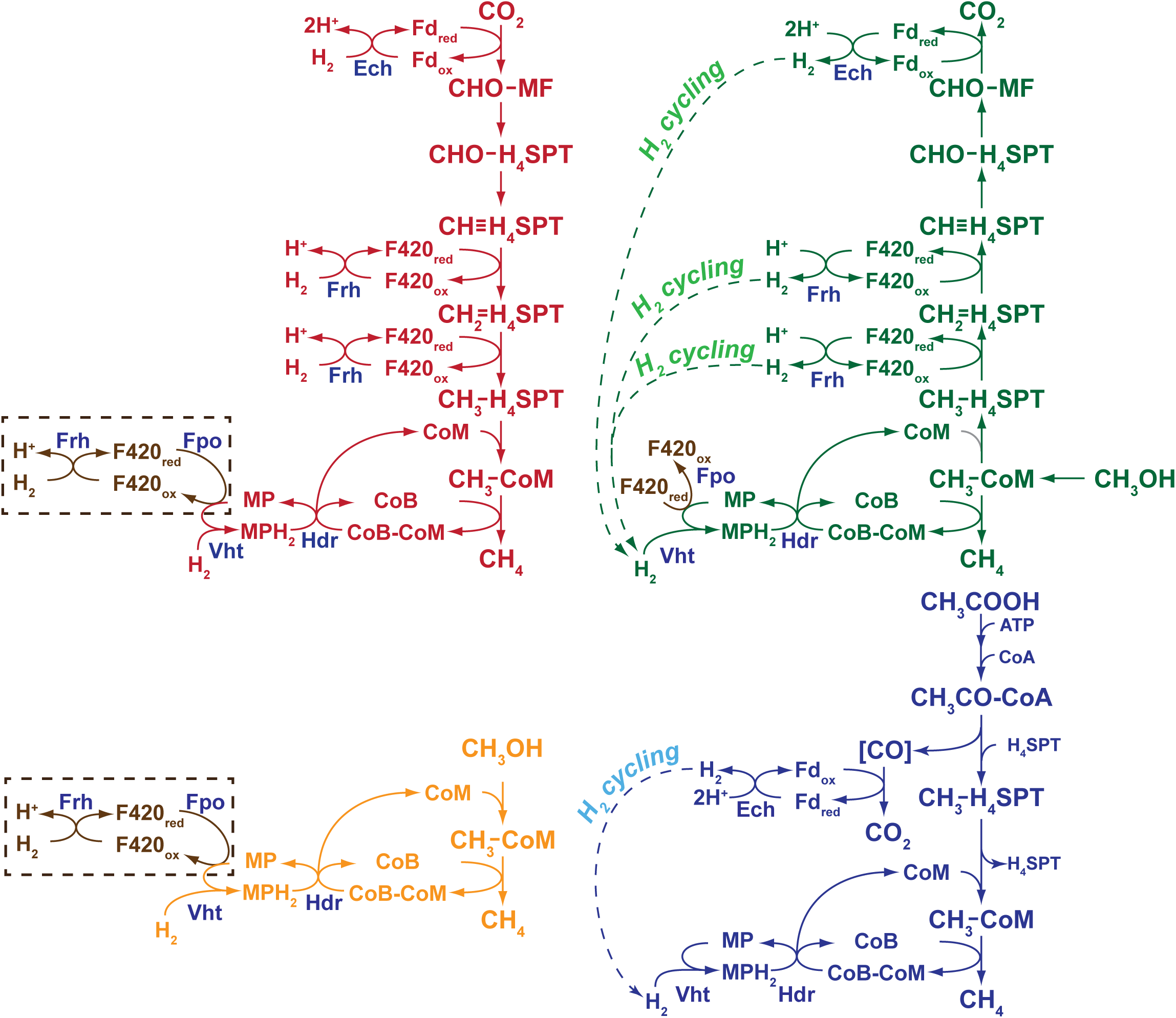
The role of H_2_ cycling in the four methanogenic pathways of *M. barkeri*. *M. barkeri* utilizes four distinct methanogenic pathways to allow growth on a variety of substrates. In the hydrogenotrophic pathway (shown in red), CO_2_ is reduced to methane using electrons derived from H_2_, while in the methyl-reducing pathway (shown in orange), H_2_ is used to reduce C-1 compounds, such as methanol, directly to CH_4_. During methylotrophic methanogenesis (shown in green), C-1 compounds are disproportionated to CO_2_ and methane, with one molecule of the C-1 compound oxidized to provide electrons for reduction of three additional molecules to methane. Finally, in the aceticlastic pathway (shown in blue), acetate is split into a methyl group and an enzyme-bound carbonyl moiety. The latter is oxidized to CO_2_ to provide electrons required for reduction of methyl group to methane. The steps catalyzed by Fpo, Frh, Vht, Ech and Hdr proteins are indicated. Steps involving H_2_ cycling are shown as labeled, hyphenated arrows. An alternate, H_2_-independent electron transport pathway is shown in brown. Experimental data support the function of this alternate pathway during methylotrophic methanogenesis, but not in hydrogenotrophic or methyl-reducing methanogenesis (as indicated by the hyphenated brown box). Abbreviations; Fpo, F420 dehydrogenase; Frh, F420-reducing hydrogenase; Vht, methanophenazine-dependent hydrogenase; Ech, energy-converting ferredoxin-dependent hydrogenase; Hdr, heterodisulfide reductase; CoM, coenzyme M; CoB, coenzyme B; CoB-CoM, mixed disulfide of CoB and CoM; MP/MPH_2_, oxidized and reduced methanophenazine; F420_ox_/F420_red_, oxidized and reduced cofactor F420; Fd_ox_/Fd_red_, oxidized and reduced ferredoxin; CHO-MF, formyl-methanofuran; H_4_SPT, tetrahydrosarcinapterin; CHO-H_4_SPT, formyl-H_4_SPT; CH≡H_4_SPT, methenyl-H_4_SPT; CH_2_=H_4_SPT, methylene-H_4_SPT; CH_3_-H_4_SPT, methyl-H_4_SPT; CH_3_-CoM, methyl-coenzyme M; CoA, coenzyme A; CH_3_CO-CoA, acetyl-coenzyme A; [CO], enzyme-bound carbonyl moiety; ATP, adenosine triphosphate.

### Depletion of Vht results in H_2_ accumulation and cell lysis

To help understand why Vht is essential, we quantified production of H_2_ and CH_4_ in cultures of the P_*tet*_*∷vht* strain with and without tetracycline (Fig 2). When grown in methanol medium in the presence of tetracycline, the accumulation of H_2_ and CH_4_ was essentially identical to that of the isogenic parent. Cultures in which *vht* is not expressed (*i.e.* without tetracycline) grew initially, but rapidly slowed, and reached an optical density that was less than half of that obtained when *vht* was expressed. The optical density subsequently dropped, suggesting cell death and lysis. Similarly, methane accumulation in cultures not expressing *vht* was much slower than in induced cultures, and only reached half of that seen under inducing conditions. In contrast, H_2_ accumulation was much higher in the absence of Vht, with final levels nearly six-fold higher than those seen in cultures that express Vht. These data clearly show that Vht is required for efficient recapture of H_2_ produced by Frh and Ech. Moreover, they suggest that H_2_ loss is responsible for the lethal consequences of *vht* repression.

### Vht is not essential in Δ*frh* mutants

If the inability to recapture H_2_ is responsible for the essentiality of Vht, then it should be possible to delete the *vht* operon in strains that do not produce hydrogen. As described above, Frh is responsible for the majority of H_2_ production during growth. Thus, we attempted to introduce a Δ*vht* allele into the Δ*frh* host. In contrast to our prior unsuccessful attempts to create a Δ*vht* single mutant, the Δ*vht*/Δ*frh* double mutant was isolated in the first attempt. Therefore, Vht is not required when Frh is absent. Like the Δ*frh* single mutant, the Δ*vht*/Δ*frh* double mutant grows slowly on methanol and produces lower levels of methane (Fig 2). Significantly, the double mutant does not produce the excessive level of H_2_ seen in the uninduced P_*tet*_*∷vht* strain, instead accumulating H_2_ at levels similar to those of the parental strain (*ca*. 20 Pa). Because Ech is the only active hydrogenase remaining in the Δ*vht*/Δ*frh* mutant, it must be responsible for H_2_ production in this strain. This begs the question of why H_2_ accumulation stops at 20 Pa in the double mutant, while the uninduced P_*tet*_*∷vht* strain produces much higher levels. We suggest that the coupling of Ech activity to generation of proton motive force thermodynamically restrains excessive H_2_ production, even in the absence of H_2_ uptake by Vht. This would also explain the viability of the Δ*vht*/Δ*frh* double mutant. This situation is in stark contrast to that seen in the *vht*-depleted strain, where the F420-dependent Frh is responsible for most of the H_2_ production (see above). Accordingly, at the low H_2_ partial pressures observed in our experiments, reduction of protons with F420 is strongly exergonic, allowing excessive hydrogen accumulation. This is also consistent with the observation that the redox state of F420 is in rapid equilibrium with H_2_ (35). Interestingly, the lower amount of H_2_ accumulation in the Δ*frh* mutant, relative to that seen in the Δ*vht/*Δ*frh* mutant, shows that Vht also consumes H_2_ produced by Ech. This supports previous studies indicating potential energy conservation via Ech/Vht H_2_ cycling during acetate metabolism (17, 23).

### *M. barkeri* has a bifurcated electron transport chain with H_2_-dependent and -independent branches

We previously showed that *M. barkeri* has a branched electron transport chain, with Frh-and F420 dehydrogenase (Fpo)-dependent branches (16). The data reported here extend our understanding of the Frh-dependent branch, and are fully consistent with the model depicted in Fig 1. Thus, during growth on methylotrophic substrates such as methanol, reduced F420 is preferentially oxidized via an energy conserving, H_2_ cycling electron transport chain that requires Frh. However, in the absence of Frh, reduced F420 is channeled into the Fpo-dependent electron transport chain, which supports growth at a significantly slower rate (Figs 2 & 4). This alternate pathway accounts for the viability of the Δ*frh* mutant, which is lost when both *frh* and *fpo* are deleted (16). Similar, but less severe phenotypes have been observed in *fpo* and *frh* mutants of *Methanosarcina mazei*, thus it seems likely that H_2_ cycling also occurs in this closely-related species (36). However, many *Methanosarcina* species, especially those that inhabit marine environments, are devoid of hydrogenase activity, despite the presence of hydrogenase encoding genes. We, and others, have interpreted this to be an adaptation to the marine environment, where H_2_-utilizing sulfate reducers are likely to disrupt H_2_ cycling due to the superior thermodynamics of H_2_ oxidation coupled to sulfate reduction (19, 37).

A similar branched electron transport chain may also explain the contradictory evidence regarding H_2_ cycling in *Desulfovibrio* species. Thus, the viability of *Desulfovibrio* hydrogenase mutants and the inability of excess H_2_ to suppress substrate catabolism can both be explained by the presence of alternative electron transport mechanisms. Indeed, metabolic modeling of *D. vulgaris* strongly supports this interpretation (13). Thus, it is critical that experiments designed to test the H_2_ cycling mechanism be interpreted within a framework that includes the possibility of branched electron transport chains. With this in mind, it seems likely that many anaerobic organisms might use H_2_ cycling for energy conservation. Indeed, since it was originally proposed, H_2_ cycling has been suggested to occur in the acetogen *Acetobacterium woodii* (10) and in the Fe (III) respiring *Geobacter sulfurreducens* (8).

### Why are Vht mutants inviable during growth on methanol/H_2_ or H_2_/CO_2_?

Although the data presented here strongly support the H_2_ cycling model, they raise additional questions regarding H_2_-dependent methanogenesis that are not easily explained. In particular, it is not readily apparent why the uninduced P_*tet*_*∷vht* mutants are inviable during hydrogenotrophic or methyl-reducing growth. As shown in Figure 4, it should be possible to channel electrons from H_2_ oxidation into the electron transport chain via Frh and Fpo. Indeed, Thauer *et al* have proposed that this alternate pathway is functional in *Methanosarcina* (38). Nevertheless, the P_*tet*_*∷vht* mutant does not grow under repressing conditions on either H_2_/CO_2_ or methanol plus H_2_. It should be stressed, that we use high concentrations of hydrogen during growth on these substrates. Thus, it is expected that reduction of F420 via Frh should be exergonic in our experiments, which would favor this pathway. (This is in contrast to the methylotrophic or aceticlastic growth conditions described above, under which the reverse reaction (*i.e.* hydrogen production) is favored.) Thus, a thermodynamic argument cannot easily explain the results. Further, based on available evidence (16, 39, 40), energy conservation via the Vht-dependent pathway should be identical to that of the alternate Frh/Fpo-dependent pathway. Thus, an energy conservation argument also cannot explain the phenomenon. One might argue that faster kinetics of the Vht-dependent pathway could be responsible, but, in our opinion, the growth (albeit slower than wild-type) of the Δ*frh* and Δ*vht/*Δ*frh* mutants during methylotrophic growth, which depends on Fpo, argues against this explanation. Therefore, as yet unknown regulatory and/or biochemical constraints on hydrogen metabolism in *Methanosarcina* await discovery.

## Materials and Methods

### Strains, media, and growth conditions

The construction and genotypes of all *Methanosarcina* strains are presented in Table S1. *Methanosarcina* strains were grown as single cells (41) at 37 °C in high salt (HS) broth medium (42) or on agar-solidified medium as described (43). Growth substrates provided were methanol (125 mM in broth medium and 50 mM in agar-solidified medium) or sodium acetate (120 mM) under a headspace of either N_2_/CO_2_ (80/20%) at 50 kPa over ambient pressure or H_2_/CO_2_ (80/20%) at 300 kPa over ambient pressure. Cultures were supplemented as indicated with 0.1% yeast extract, 0.1% casamino acids, 10 mM sodium acetate or 10 mM pyruvate. Puromycin (CalBioChem, San Diego, CA) was added at 2 μg/ml for selection of the puromycin transacetylase (*pac*) gene (33). 8-aza-2,6-diaminopurine (8-ADP) (Sigma, St Louis, MO) was added at 20 μg/ml for selection against the presence of *hpt* (33). Tetracycline was added at 100 μg/ml to induce the tetracycline-regulated P*mcrB*(*tet*O3) promoter (34). Standard conditions were used for growth of *Escherichia coli* strains (44) DH5α/λ-*pir* (45) and DH10B (Stratagene, La Jolla, CA), which were used as hosts for plasmid constructions.

### DNA methods and plasmid construction

Standard methods were used for plasmid DNA isolation and manipulation using *E. coli* hosts (46). Liposome mediated transformation was used for *Methanosarcina* as described (47). Genomic DNA isolation and DNA hybridization were as described (32, 42, 43). DNA sequences were determined from double-stranded templates by the W.M. Keck Center for Comparative and Functional Genomics, University of Illinois. Plasmid constructions are described in the supporting information (Tables S2 & S3).

### Construction of the Δ*frh* and Δ*vht/*Δ*frh* mutants

The markerless genetic exchange method (33) using plasmid pGK4 was employed to delete *frhADGB* (Δ*frh*) in the Δ*hpt* background of *M. barkeri* Fusaro (Tables S1, S2, & S3) using methanol/H_2_/CO_2_ as the growth substrate. The Δ*vht/*Δ*frh* mutant was constructed by deleting *vhtGACD* in the Δ*frh* markerless mutant by the homologous recombination-mediated gene replacement method (32). To do this, the 5.6 kb XhoI/NotI fragment of pGK82B was used to transform the Δ*frh* mutant to puromycin resistance on methanol medium. The mutants were confirmed by PCR and DNA hybridization (data not shown).

### Construction of the tetracycline-regulated *vht* mutant (P_*tet*_*∷vht*)

The tetracycline-regulated P*mcrB*(*tet*O3) promoter was employed to drive conditional expression of the *vht* operon in *M. barkeri* WWM157 (34). This strain was constructed by transforming WWM154 to puromycin reistance using the 7 kb NcoI/SpeI fragment of pGK61A (Tables S1, S2, & S3). The transformants were selected on methanol plus H_2_/CO_2_ medium in the presence of puromycin and tetracycline. The P_*tet*_*∷vht* strain was confirmed by DNA hybridization (data not shown). To ensure that the native *vht* promoter (P*vht*) did not interfere with expression from P*mcrB*(*tet*O3), 382 bp upstream of *vhtG* were deleted in P_*tet*_*∷vht*. This left 1038 bp intact for the expression of the *hyp* operon, which is upstream of the *vht* operon and expressed in the opposite direction.

### Determination of Vht essentiality during growth on all substrate-types

Growth of WWM157 (P_*tet*_*∷vht*) and WWM154 (isogenic parent) on methanol, methanol/H_2_/CO_2_, H_2_/CO_2_, and acetate were analyzed by the spot-plate method (48). Cultures were first adapted for at least 15 generations to the substrate of interest; tetracycline was added to each medium for growth of WWM157. Upon reaching stationary phase, 10 ml of culture was washed three times and re-suspended in 5 ml HS medium that lacked growth substrate. Subsequently, 10 µl of 10-fold serial dilutions was spotted onto the following: 3 layers of GB004 paper (Whatman, NJ), 2 layers of GB002 paper (Schleicher & Schuell BioScience, NH), 1 layer of 3MM paper (Whatman, NJ), and a 0.22 mM nylon membrane (GE Water and Process Technologies, PA) soaked in 43 ml of HS-medium containing the substrate of interest with and without Tc. Plates were sealed and incubated at 37 °C for at least two weeks in an intrachamber anoxic incubator (49). Growth on acetate and methanol was tested under an atmosphere of N_2_/CO_2_/H_2_S (80/19.9/0.1 ratio), while growth on methanol/H_2_/CO_2_ or H_2_/CO_2_ was tested under an atmosphere of H_2_/CO_2_/H_2_S (80/19.9/0.1 ratio).

### Measurement of H_2_, CH_4_ and OD_600_ during growth on methanol

*M. barkeri* WWM85 (isogenic parent), WWM157 (P_*tet*_*∷vht;* grown in presence of Tc), WWM115 (Δ*frh*) and WWM351 (Δ*vht/*Δ*frh*) were grown on methanol until mid-exponential phase (OD_600_ *c.* 0.5) and then 1 ml (WWM85 and WWM157) or 5ml (WWM115 and WWM351) were inoculated into 100 ml HS-methanol in a 500 ml serum bottle. For WWM157, the culture was washed once prior to inoculation with or without tetracycline. To measure H_2_ and CH_4_, ca. 1 ml or 2 ml headspace sample was withdrawn aseptically from the culture at various time points with a syringe that had been flushed with sterile, anaerobic N_2_. The gas sample was then diluted into 70 ml helium. A gas-tight syringe flushed with helium was subsequently used to withdraw 3 ml of the diluted sample, which was then injected into an SRI gas chromatograph, equipped with a reduction gas detector (RGD) and a thermal conductivity detector (TCD) at 52°C. The RGD column was a three-foot long 13X molecule sieve, whereas the TCD column was a six-foot HayeSep D. RGD was used to detect H_2_ by peak height and TCD for CH_4_ by peak area. Helium was used as the carrier gas. OD_600_ was also measured during the growth curve.

## Acknowledgements

We wish to thank Rob Sanford for providing assistance and facilities for measurement of low hydrogen partial pressures. The authors acknowledge the Division of Chemical Sciences, Geosciences, and Biosciences, Office of Basic Energy Sciences of the U.S. Department of Energy through Grant DE-FG02-02ER15296 for funding of this work.

---

**Figure S1. Hydrogenase operons in *Methanosarcina barkeri*.** Three distinct types of hydrogenase are encoded by *M. barkeri* Fusaro. The *frh* and *fre* operons encode putative F420-reducing hydrogenases, while the *vht* and *vhx* operons encode putative methanophenazine-reducing hydrogenases. Genetic and biochemical data show that neither the *fre* nor the *vhx* operon is capable of producing an active hydrogenase under any growth condition yet examined (T.D. Mand, G. Kulkarni, and W.W. Metcalf, 2018, bioRxiv doi: https://doi.org/10.1101/334656). The *ech* operon encodes a ferredoxin-dependent energy-conserving hydrogenase. The locus tags are shown below each gene, with the prefix “Mbar_” omitted to save space (indicated by an asterisk) in some cases.

## References

1. White D. 2000. Electron Transport, p 103–131, The Physiology and Biochemistry of Prokaryotes, 2nd ed. Oxford University Press, New York.

2. White D. 2000. Bioenergetics in the Cytosol, p 165–179, The Physiology and Biochemistry of Prokaryotes, 2nd ed. Oxford University Press, New York.

3. Gottschalk G, Thauer RK. 2001. The Na^+^-translocating methyltransferase complex from methanogenic archaea. Biochim Biophys Acta 1505:28–36.

4. Anantharam V, Allison MJ, Maloney PC. 1989. Oxalate:formate exchange. The basis for energy coupling in *Oxalobacter*. J Biol Chem 264:7244–7250.

5. Kuhner CH, Hartman PA, Allison MJ. 1996. Generation of a proton motive force by the anaerobic oxalate-degrading bacterium *Oxalobacter formigenes*. Appl Environ Microbiol 62:2494–2500.

6. Béjà O, Aravind L, Koonin EV, Suzuki MT, Hadd A, Nguyen LP, Jovanovich SB, Gates CM, Feldman RA, Spudich JL, Spudich EN, DeLong EF. 2000. Bacterial rhodopsin: evidence for a new type of phototrophy in the sea. Science 289:1902– 1906.

7. Odom JM, Peck HD. 1981. Hydrogen cycling as a general mechanism for energy coupling in the sulfate-reducing bacteria, *Desulfovibrio* sp. FEMS Microbiol Lett 12:47–50.

8. Coppi MV. 2005. The hydrogenases of *Geobacter sulfurreducens*: a comparative genomic perspective. Microbiology 151:1239–1254.

9. Lovley DR, Ferry JG. 1985. Production and Consumption of H_2_ during Growth of *Methanosarcina* spp. on Acetate. Appl Environ Microbiol 49:247–249.

10. Odom JM, Peck HD. 1984. Hydrogenase, electron-transfer proteins, and energy coupling in the sulfate-reducing bacteria *Desulfovibrio*. Annu Rev Microbiol 38:551–592.

11. Lupa B, Hendrickson EL, Leigh JA, Whitman WB. 2008. Formate-dependent H_2_ production by the mesophilic methanogen *Methanococcus maripaludis*. Appl Environ Microbiol 74:6584–6590.

12. Peck HD, Jr., LeGall J, Lespinat PA, Berlier Y, Fauque G. 1987. A direct demonstration of hydrogen cycling by *Desulfovibrio vulgaris* employing membrane-inlet mass spectrometry. FEMS Microbiol Lett 40:295–299.

13. Noguera DR, Brusseau GA, Rittmann BE, Stahl DA. 1998. A unified model describing the role of hydrogen in the growth of *Desulfovibrio vulgaris* under different environmental conditions. Biotechnol Bioeng 59:732–746.

14. Odom JM, Wall JD. 1987. Properties of a hydrogen-inhibited mutant of *Desulfovibrio desulfuricans* ATCC 27774. J Bacteriol 169:1335–1337.

15. Pankhania IP, Gow LA, Hamilton WA. 1986. The Effect of Hydrogen on the Growth of Desulfovibrio vulgaris (Hildenborough) on Lactate. Journal of General Microbiology 132:3349–3356.

16. Kulkarni G, Kridelbaugh DM, Guss AM, Metcalf WW. 2009. Hydrogen is a preferred intermediate in the energy-conserving electron transport chain of *Methanosarcina barkeri*. Proc Natl Acad Sci U S A 106:15915–15920.

17. Meuer J, Kuettner HC, Zhang JK, Hedderich R, Metcalf WW. 2002. Genetic analysis of the archaeon *Methanosarcina barkeri* Fusaro reveals a central role for Ech hydrogenase and ferredoxin in methanogenesis and carbon fixation. Proc Natl Acad Sci U S A 99:5632–5637.

18. Deppenmeier U, Blaut M, Lentes S, Herzberg C, Gottschalk G. 1995. Analysis of the *vhoGAC* and *vhtGAC* operons from *Methanosarcina mazei* strain Gö1, both encoding a membrane-bound hydrogenase and a cytochrome *b*. Eur J Biochem 227:261–269.

19. Guss AM, Kulkarni G, Metcalf WW. 2009. Differences in hydrogenase gene expression between *Methanosarcina acetivorans* and *Methanosarcina barkeri*. J Bacteriol 191:2826–2833.

20. Vaupel M, Thauer RK. 1998. Two F_420_-reducing hydrogenases in *Methanosarcina barkeri*. Arch Microbiol 169:201–205.

21. Brodersen J, Bäumer S, Abken HJ, Gottschalk G, Deppenmeier U. 1999. Inhibition of membrane-bound electron transport of the methanogenic archaeon *Methanosarcina mazei* Gö1 by diphenyleneiodonium. Eur J Biochem 259:218– 224.

22. Mand TD, Kulkarni G, Metcalf WW. 2018. Genetic, biochemical, and molecular characterization of *Methanosarcina barkeri* mutants lacking three distinct classes of hydrogenase. bioRxiv doi: https://doi.org/10.1101/334656

23. Meuer J, Bartoschek S, Koch J, Künkel A, Hedderich R. 1999. Purification and catalytic properties of Ech hydrogenase from *Methanosarcina barkeri*. Eur J Biochem 265:325–335.

24. Peters JW, Schut GJ, Boyd ES, Mulder DW, Shepard EM, Broderick JB, King PW, Adams MWW. 2015. [FeFe]-and [NiFe]-hydrogenase diversity, mechanism, and maturation. Biochim Biophys Acta 1853:1350–1369.

25. Phelps TJ, Conrad R, Zeikus JG. 1985. Sulfate-Dependent Interspecies H_2_ Transfer between *Methanosarcina barkeri* and *Desulfovibrio vulgaris* during Coculture Metabolism of Acetate or Methanol. Appl Environ Microbiol 50:589–594.

26. Bhatnagar L, Krzycki JA, Zeikus JG. 1987. Analysis of hydrogen metabolism in *Methanosarcina barkeri*: Regulation of hydrogenase and role of CO-dehydrogenase in H_2_ production. FEMS Microbiol Lett 41:337–343.

27. Boone DR, Menaia JAGF, Boone JE, Mah RA. 1987. Effects of Hydrogen Pressure during Growth and Effects of Pregrowth with Hydrogen on Acetate Degradation by *Methanosarcina* Species. Appl Environ Microbiol 53:83–87.

28. Krzycki JA, Morgan JB, Conrad R, Zeikus JG. 1987. Hydrogen metabolism during methanogenesis from acetate by *Methanosarcina barkeri*. FEMS Microbiol Lett 40:193–198.

29. Ahring BK, Westermann P, Mah RA. 1991. Hydrogen inhibition of acetate metabolism and kinetics of hydrogen consumption by *Methanosarcina thermophila* TM-1. Arch Microbiol 157:38–42.

30. Zinder SH, Anguish T. 1992. Carbon Monoxide, Hydrogen, and Formate Metabolism during Methanogenesis from Acetate by Thermophilic Cultures of *Methanosarcina* and *Methanothrix* Strains. Appl Environ Microbiol 58:3323–3329.

31. Maeder DL, Anderson I, Brettin TS, Bruce DC, Gilna P, Han CS, Lapidus A, Metcalf WW, Saunders E, Tapia R, Sowers KR. 2006. The *Methanosarcina barkeri* genome: comparative analysis with *Methanosarcina acetivorans* and *Methanosarcina mazei* reveals extensive rearrangement within methanosarcinal genomes. J Bacteriol 188:7922–7931.

32. Zhang JK, White AK, Kuettner HC, Boccazzi P, Metcalf WW. 2002. Directed mutagenesis and plasmid-based complementation in the methanogenic archaeon *Methanosarcina acetivorans* C2A demonstrated by genetic analysis of proline biosynthesis. J Bacteriol 184:1449–1454.

33. Pritchett MA, Zhang JK, Metcalf WW. 2004. Development of a markerless genetic exchange method for *Methanosarcina acetivorans* C2A and its use in construction of new genetic tools for methanogenic archaea. Appl Environ Microbiol 70:1425–1433.

34. Guss AM, Rother M, Zhang JK, Kulkarni G, Metcalf WW. 2008. New methods for tightly regulated gene expression and highly efficient chromosomal integration of cloned genes for *Methanosarcina* species. Archaea 2:193–203.

35. de Poorter LM, Geerts WJ, Keltjens JT. 2005. Hydrogen concentrations in methane-forming cells probed by the ratios of reduced and oxidized coenzyme F_420_. Microbiology 151:1697–1705.

36. Welte C, Deppenmeier U. 2011. Re-evaluation of the function of the F_420_ dehydrogenase in electron transport of *Methanosarcina mazei*. FEBS J 278:1277–1287.

37. Deppenmeier U. 2004. The membrane-bound electron transport system of *Methanosarcina* species. J Bioenerg Biomembr 36:55–64.

38. Thauer RK, Kaster A-K, Seedorf H, Buckel W, Hedderich R. 2008. Methanogenic archaea: ecologically relevant differences in energy conservation. Nat Rev Microbiol 6:579–591.

39. Ide T, Baumer S, Deppenmeier U. 1999. Energy conservation by the H_2_:heterodisulfide oxidoreductase from *Methanosarcina mazei* Gö1: identification of two proton-translocating segments. J Bacteriol 181:4076–4080.

40. Baumer S, Ide T, Jacobi C, Johann A, Gottschalk G, Deppenmeier U. 2000. The F_420_H_2_ dehydrogenase from *Methanosarcina mazei* is a Redox-driven proton pump closely related to NADH dehydrogenases. J Biol Chem 275:17968–17973.

41. Sowers KR, Boone JE, Gunsalus RP. 1993. Disaggregation of *Methanosarcina* spp. and Growth as Single Cells at Elevated Osmolarity. Appl Environ Microbiol 59:3832–3839.

42. Metcalf WW, Zhang JK, Shi X, Wolfe RS. 1996. Molecular, genetic, and biochemical characterization of the *serC* gene of *Methanosarcina barkeri* Fusaro. J Bacteriol 178:5797–5802.

43. Boccazzi P, Zhang JK, Metcalf WW. 2000. Generation of dominant selectable markers for resistance to pseudomonic acid by cloning and mutagenesis of the *ileS* gene from the archaeon *Methanosarcina barkeri* Fusaro. J Bacteriol 182:2611–2618.

44. Wanner BL. 1986. Novel regulatory mutants of the phosphate regulon in *Escherichia coli* K-12. J Mol Biol 191:39–58.

45. Miller VL, Mekalanos JJ. 1988. A novel suicide vector and its use in construction of insertion mutations: osmoregulation of outer membrane proteins and virulence determinants in *Vibrio cholerae* requires toxR. J Bacteriol 170:2575–2583.

46. Ausubel FM, Brent R, Kingston RE, Moore DD, Seidman JG, Smith JA, Struhl K. 1992. Current Protocols in Molecular Biology. Wiley & Sons, New York.

47. Metcalf WW, Zhang JK, Apolinario E, Sowers KR, Wolfe RS. 1997. A genetic system for Archaea of the genus *Methanosarcina*: liposome-mediated transformation and construction of shuttle vectors. Proc Natl Acad Sci U S A 94:2626–2631.

48. Buan N, Kulkarni G, Metcalf W. 2011. Genetic methods for *Methanosarcina* species. Methods Enzymol 494:23–42.

49. Metcalf WW, Zhang JK, Wolfe RS. 1998. An anaerobic, intrachamber incubator for growth of *Methanosarcina* spp. on methanol-containing solid media. Appl Environ Microbiol 64:768–770.

